# Cognitive neuropsychological and neuroanatomic predictors of naturalistic action performance in left hemisphere stroke: a retrospective analysis

**DOI:** 10.1101/2024.07.01.601398

**Authors:** Simon Thibault, Aaron L. Wong, Laurel J. Buxbaum

## Abstract

Individuals who have experienced a left hemisphere cerebrovascular accident (LCVA) have been shown to make errors in naturalistic action tasks designed to assess the ability to perform everyday activities such as preparing a cup of coffee. Naturalistic action errors in this population are often attributed to limb apraxia, a common deficit in the representation and performance of object-related actions. However, naturalistic action impairments are also observed in right hemisphere stroke and traumatic brain injury, populations infrequently associated with apraxia, and errors across all these populations are influenced by overall severity. Based on these and other data, an alternative (though not mutually exclusive) account is that naturalistic action errors in individuals with LCVA are also a consequence of deficits in general attentional resource availability or allocation. In this study, we conducted a retrospective analysis of data from a group of 51 individuals with LCVA who had completed a test of naturalistic action, along with a battery of tests assessing praxis, attention allocation and control, reasoning, and language abilities to determine which of these capacities contribute uniquely to naturalistic action impairments. Using a regularized regression method, we found that naturalistic action impairments are predicted by both praxis deficits (hand posture sequencing and gesture recognition), as well as attention allocation and control deficits (orienting and dividing attention), along with language comprehension ability and age. Using support vector regression-lesion symptom mapping, we demonstrated that naturalistic action impairments are associated with lesions to posterior middle temporal gyrus and anterior inferior parietal lobule regions known to be implicated in praxis; as well the middle frontal gyrus that has been implicated in both praxis and attention allocation and control. Together, these findings support the hypothesis that naturalistic action impairments in people with LCVA are a consequence of apraxia as well as deficits in attention allocation and control.

## Introduction

Naturalistic actions such as preparing toast with butter and jam require the use of different manipulable objects (e.g. knife, spoon, toaster). Each object in turn requires knowledge about how it should be used. This manipulation knowledge is known to be subserved by a so-called *praxis system* that enables the production and recognition of tool-related actions (e.g., pantomiming how a tool is to be used) and the imitation of both meaningful and meaningless actions (Binkofski & Buxbaum, 2013; Buxbaum & Kalénine, 2010, 2022). The praxis system is primarily supported by the left hemisphere (e.g., Binkofski and Buxbaum, 2013; Buxbaum and Kalénine, 2010; Johnson-Frey, 2004; Lewis, 2006; Orban and Caruana, 2014). It is therefore perhaps unsurprising that people who have experienced a left hemisphere cerebrovascular accident (LCVA) make errors in naturalistic action tasks at rates higher than neurotypicals (Buxbaum et al., 1998). Such errors include omitting a step in the task (e.g., failing to put butter on toast) or inverting the appropriate order of task steps (e.g., spreading butter on the bread before toasting it; Buxbaum et al., 1998). It has also been demonstrated that gesture pantomime, a task commonly used to assess praxis function, correlates with performance of naturalistic actions in individuals with LCVA (Bienkiewicz et al., 2015; Hartmann et al., 2005; Poole et al., 2011) and that the use of familiar tools correlates with a breakfast preparation task (Buchmann & Randerath, 2017).

However, naturalistic action impairments have also been observed in individuals less likely to have praxis deficits such as those who have suffered a right cerebral vascular accident (RCVA; Bickerton et al., 2012, 2007; Bienkiewicz et al., 2015; Hartmann et al., 2005), traumatic brain injury (TBI; Buxbaum et al., 1998; Schwartz et al., 1998), or Parkinson’s disease (Balouch & Rusted, 2014; Giovannetti et al., 2002, 2008, 2012; Roll et al., 2017). Moreover, patterns of error types are highly similar across these populations (Buxbaum et al., 1998; Giovannetti et al., 2002; Jarry et al., 2021; Schwartz et al., 1998, 1999). These data suggest that naturalistic action impairments may arise for reasons additional to deficits of the left-lateralized praxis system. Based in part on evidence that dividing attention with a secondary task disrupts performance of the primary naturalistic action task (Giovannetti et al., 2007; Humphreys & Riddoch, 2000), the limited-capacity resources theory proposes that errors in naturalistic tasks may instead arise because of a reduction in general attentional resources after damage to the brain (Giovannetti et al., 2007; Reason, 1984; Robertson et al., 1997; Schwartz et al., 1998). It has been suggested that limited attentional resources may influence the naturalistic action abilities of people with RCVA, whereas underlying deficits in lexical-semantic knowledge may be relatively more influential in people with LCVA (Rumiati, 2005). On this account, it is unclear whether attentional resource limitations affect naturalistic task performance even in LCVA and suggest the need for additional investigation. Research on attention distinguishes between attention control and attention allocation. Attention control (also termed cognitive control, executive control, or executive attention, e.g., Burgoyne and Engle, 2020) is the ability to select relevant thoughts and behaviors and suppress inappropriate thoughts and behaviors based on current goals and task demands (Miller & Cohen, 2001). Attention control is particularly relevant in situations of increased cognitive demand, such as during distraction. Attention allocation (sometimes referred to as “alerting”), a lower-level function that does not depend on volition (Anderson, 2021), is the ability to replace a resting state with a state of preparation to detect and respond to inputs (Petersen & Posner, 2012). These two different processes are highly interdependent and provide the foundation for other cognitive functions such as working memory and decision-making (Boshra & Kastner, 2022). From a neuroanatomic perspective, attention allocation and control are known to rely on a bilateral brain network involving fronto-temporo-parietal areas, midline structures such as the anterior cingulate, and subcortical connections with the brainstem and thalamus (Boshra & Kastner, 2022; Fan et al., 2005). The bilateral involvement of the brain in attention allocation and control may explain why these processes are impaired in numerous neurological conditions (Manohar et al., 2014). However, whether such deficits contribute to naturalistic action impairments remains a matter of debate. Indeed, most of the evidence in dementia populations has not distinguished between specific attention allocation/control deficits and overall disease severity (Giovannetti et al., 2002; Jarry et al., 2021).

In this study, we assessed whether naturalistic action impairments in people with LCVA arise from a combination of praxis and attention allocation/control deficits using a retrospective analysis of data from 51 people with chronic LCVA. The extensive dataset included performance on a test of naturalistic action along with 20 predictors derived from a battery of tests measuring praxis, attention processes, reasoning, and language; we also included participant demographics. We tested which predictors accounted for naturalistic action impairments. We also assessed whether performance on the test of naturalistic action was related to individuals’ reports of difficulties in daily activities. Finally, using a machine-learning based approach, we performed an exploratory analysis of the lesion correlates of naturalistic action impairments.

## Material and Methods

### Participants

Data for the study were collected in 2003-2006 but remained unpublished. Community-dwelling participants with chronic left hemisphere stroke were recruited from the Research Registry of Moss Rehabilitation Research Institute and from outpatient admissions at MossRehab. All procedures were conducted at Moss Rehabilitation Research Institute. Participants were between the ages of 18 and 80, had sustained a single left-hemisphere cerebrovascular accident (LCVA) more than six months prior to participation, and did not have a history of major psychiatric diagnosis, dementia, or current drug/alcohol abuse. Informed consent was obtained for all participants in accordance with the policies of the Institutional Review Board of Einstein Healthcare Network. Participants were assessed to ensure that they exhibited at least some residual neurological impairment as determined by a score greater than 0 on the National Institutes of Health (NIH) stroke scale (Brott et al., 1989), but did not have severe language comprehension deficits as determined by a score of four or more on the comprehension subscale of the Western Aphasia Battery (WAB; Kertesz and Hooper, 1982). This retrospective dataset included a total of 64 participants who performed the main task (Naturalistic Action Test – NAT; Schwartz et al., 2002). Of those individuals, 12 individuals did not complete one or more of the relevant tests from the behavioral test battery. Visual inspection of research-quality scans conducted after behavioral testing revealed that one participant had a lesion in the right hemisphere; thus, this participant was excluded. The remaining 51 participants were included in our analyses. Demographic information for these participants is shown in Table 1. Data of four left-handed participants were collected and included in the model with the other right-handed participants. This choice was made based on previous studies that show similar praxis impairments in brain damaged left and right handers (Borod et al., 1985; Goldenberg, 2013; but see Kroliczak et al., 2021). Inspection of the data did not reveal obvious differences in performance between left and right handers.

**Table 1:**
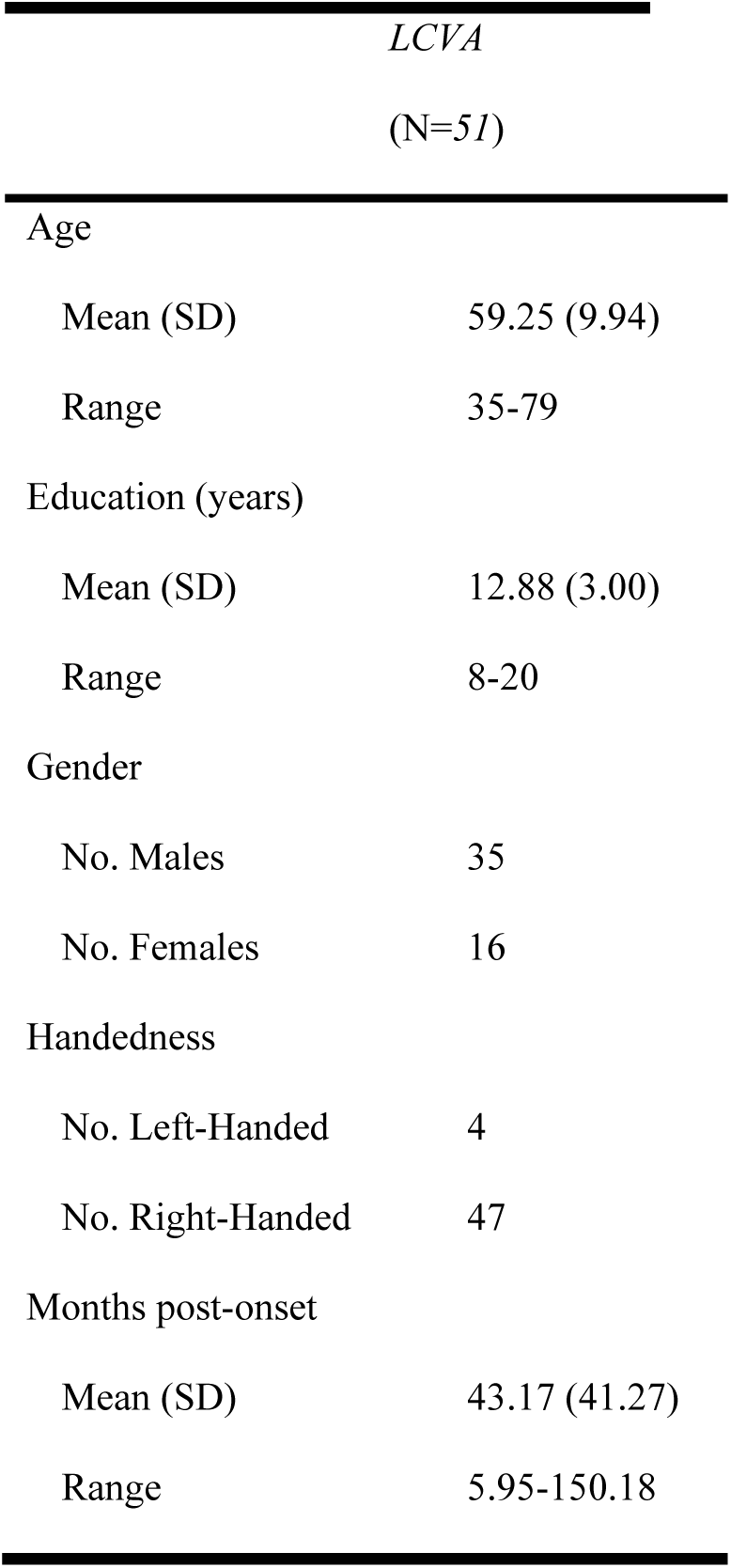
Demographic information.

### Naturalistic Action Test (NAT)

The NAT (Schwartz et al., 2002) quantifies performance during three naturalistic tasks of increasing complexity: preparing a cup of coffee and toast with butter and jam, wrapping a gift, and preparing a lunchbox and packing a schoolbag (Table 2). To ensure comprehension, instructions were read aloud to participants while they viewed black-and-white drawings of the completed task (e.g., a slice of toast with butter and jam). Following instructions, the examiner asked if the participant understood what he/she was expected to do; if not, the instructions were repeated until the participant indicated that he/she understood them. The instructions also included a familiarization phase where the participants had to point to the object named by the experimenter to ensure that they recognized each object before starting the task. Physical assistance with bimanual aspects of the task (e.g., stabilization of objects) was provided for individuals who required it. According to a detailed scoring protocol (Schwartz et al., 2002), participants received a total NAT score from zero to 18 (zero to six on each of the three tasks). A score of 18 indicated perfect performance. After informed consent was obtained, the NAT was administered during an initial session that also included the Western Aphasia Battery and NIH stroke scale. This session required approximately 1.5 hours to complete.

**Table 2:** Overview of the Naturalistic Action Test.

| *Item* |  | *Objects arrayed on table* | *Additional requirements* |
| --- | --- | --- | --- |
| 1 | Make toast with butter and jelly and instant coffee with cream and sugar | Target objects | None |
| 2 | Wrap a gift as a present | Target objects and distractors* | None |
| 3 | Prepare and pack a child's lunchbox (with sandwich, drink, and cookies); pack a child's schoolbag (with notebook and stocked pencil case) | Target objects and closed drawer containing more targets plus distractors** | Signal end of each task by activating bell attached to underside of tabletop |

\*Electric tape, stapler, paper bag, gardening shears; \*\*Contents of drawer: (targets) knife, thermos lid, thermos cup; (distractors) envelope, spoon, fork, extra knife, coupons, spatula, retractable tape measure, spool of thread, screwdriver, toothbrush, and tongs.

### Behavioral Test Battery

To examine predictors of NAT performance, participants also completed a battery of tests assessing praxis, attention allocation and control, reasoning, and language. The tests are described below and were administered over three separate testing sessions within two weeks (4-6 hours of testing for each subject). The orders of the sessions, as well as the order of tasks within each session were randomized using a Latin Square design. The sample performance in each test is reported in Table 3. Because praxis is typically assessed with the hand ipsilesional to the stroke to avoid confounds due to motor impairments, participants used their less-impaired left hand in the praxis tasks (Hand Posture Sequencing, Gesture to Sight of Object, and Gesture Imitation). All other tasks allowed participants to respond with their preferred hand.

**Table 3:** Summary of data from the Naturalistic Action Test, predictor tasks assessing praxis, attention allocation and control, reasoning, and language, as well as demographic variables and measures of functional independence.

| **Task** | **Mean** | **Standard Deviation** | **Range** |
| --- | --- | --- | --- |
| Naturalistic Action Test (Points) | 15 | 3.38 | 3 - 18 |
| **Praxis** |  |  |  |
| Function Similarity (% Correct) | 82.96 | 8.72 | 58 - 96 |
| Manipulation Similarity (% Correct) | 83.46 | 10.87 | 48 - 100 |
| GR Semantics (Number Correct) | 22.33 | 1.93 | 15 - 24 |
| GR Spatial (Number Correct) | 20.17 | 2.82 | 8 - 24 |
| Gesture to Sight of Object (% Correct) | 84.30 | 9.86 | 55 - 97 |
| Novel Tool Use (Points) | 22.02 | 2.52 | 12 - 24 |
| Hand Posture Sequencing Task (Number Errors) | 7.02 | 9.35 | 0 - 48 |
| Meaningless Imitation (Number Correct) | 45.81 | 5.80 | 33 - 56 |
| **Attention control and allocation** |  |  |  |
| ANT alerting (Normalized Reaction Time) | 0.04 | 0.05 | -0.09 - 0.16 |
| ANT orienting (Normalized Reaction Time) | 0.03 | 0.06 | -0.01 - 0.17 |
| ANT accuracy (% Correct) | 89.21 | 12.38 | 58.68 - 100 |
| Digit Span (Number Correct) | 4.15 | 1.61 | 1 - 7 |
| Dual-Task Interference (Reaction Time, ms.) | 768 | 993 | 44 - 3996 |
| Self-Ordered Pointing Test (Number Correct) | 8.08 | 1.94 | 4 - 12 |
| Sustained Attention to Response Test (d-prime) | 2.72 | 1.16 | -0.39 - 4.86 |
| **Reasoning** |  |  |  |
| Tower of London (Number Correct) | 7.98 | 1.91 | 4 - 12 |
| Brixton (Number Correct) | 22.16 | 8.01 | 9 - 48 |
| **Language** |  |  |  |
| WAB Comprehension (Points) | 9.12 | 1.09 | 6.05 - 10 |
| **Demographics** |  |  |  |
| Age (Years) | 59.54 | 10.13 | 35 - 79 |
| Months Post Stroke onset (Months) | 44.01 | 41.82 | 5.95 - 150.18 |
| **Functional Independence** |  |  |  |
| OARS-MFAQ (Points) | 10.61 | 2.64 | 4 - 14 |

### Praxis Tasks

#### Function and Manipulation Similarity Test

The function and manipulation similarity test measured the ability to access knowledge about the purpose and actions associated with tools. Each test item consisted of a pair of black-and-white drawings of common manipulable objects presented using Psyscope (Boronat et al., 2005). Participants were asked to decide whether the two depicted objects had the same function (Function condition; e.g., air conditioner and fan) or the same manner of manipulation (Manipulation condition; e.g., computer keyboard and piano). Participants were scored on the correct proportion of responses on the 64 items in each condition.

#### Gesture recognition

The gesture recognition test measured the ability to recognize the actions associated with common tools. The gesture recognition test was closely based on a test developed previously (Gonzalez Rothi et al., 1985; Heilman et al., 1982; see Tarhan et al., 2015). On each trial, participants saw the printed name of a gesture (e.g., “hammering”) for 5000 ms and heard the name of the gesture spoken twice. Participants then watched two sequential videos of object-related gestures being pantomimed and had to decide which of the two viewed gestures matched the named gesture. In the Semantic condition (24 items), one of the gestures in the pair was incorrect by virtue of a semantic relationship to the target gesture (e.g., sawing). In the Spatiotemporal condition (24 items), one of the gestures in the pair was incorrect by virtue of a spatial error in hand posture (eight items), arm posture/trajectory (eight items), or amplitude and/or timing (eight items). The conditions were blocked, and the order of correct and incorrect videos on each trial was randomized. Performance was scored as the number of gestures correctly recognized in each condition.

#### Gesture to Sight of Object

The gesture to sight of object test, a classic measure of praxis, assessed the retrieval and execution of tool-use actions. Participants were shown an object on each of 10 trials and were asked to pantomime its use using their left hand (Buxbaum et al., 2000, 2005). Participants were not allowed to touch or manipulate the object. Scoring was analogous to that of the meaningless gesture imitation task, with the addition of a “content” category. Content was scored as zero if another recognizable gesture was substituted for the target gesture (e.g., brushing teeth when the object was a comb) and one otherwise. Test items that received a content score of zero were not scored on the other four spatiotemporal components. Using a detailed scoring protocol performed by trained coders with established inter-rater reliability (Cohen’s Kappa > 85%), the percentage of correct gestures was calculated across items that received a content score of one (Buxbaum et al., 2000).

#### Hand Posture Sequencing Task

The hand posture sequencing task measured the ability to execute a sequence of gestures based upon a visual cue. Participants were asked to produce 20 sequences of five hand postures each using the left hand (task modeled after Harrington and Haaland, 1991). Each hand posture sequence comprised some combination of handle grasping, button pressing, and palming a computer mouse in non-repeating order, as indicated by a series of five symbols displayed on a computer screen. The symbols were semi-abstract in that the symbol for palming was a solid square, the symbol for button press was a dot within a circle, and the symbol for grasping the handle was a handle-shaped “U”. The task was administered using Psyscope on a Macintosh computer. Before beginning the task, participants were trained across 45 trials (15 trials with each posture) to correctly identify the symbols associated with each action in the sequence. In the second phase of practice, the participants executed ten sequences like the ones presented for the main task. An additional two rounds of ten trials of practice (20 additional trials total) were conducted if necessary. The task was stopped if participants could not reliably execute each posture in response to the presented symbol by the end of the practice phase. Each sequence trial with the ipsilesional left hand resting on the space bar of a computer keyboard while viewing a fixation cross on the monitor. The cross brightened to alert participants to an upcoming trial. A series of five symbols was displayed to denote the sequence of postures to be made. After a jittered delay (1000-2000 ms), a tone signaled participants to lift the hand and produce the series of movements on the apparatus, which contained a row of five handles, a row of five doorbells, and a row of five computer mice. The symbols remained on the monitor during the response. Responses were videotaped; accuracy was scored offline using these videos. Errors included 1) taking more than 2000 ms to initiate a movement during the RT interval, 2) executing an incorrect hand posture, or 3) executing an incorrect hand posture before correction (i.e., corrected errors). These three error sub-types were summed to produce a total error score.

#### Meaningless Gesture Imitation

The meaningless gesture imitation test, a classic measure of praxis, assessed the ability to produce novel gestures demonstrated by a model. Participants were asked to copy gestures performed by a videotaped model as precisely as possible, using their less impaired (ipsilesional) left hand (Goldenberg & Hagmann, 1997). The model used their right hand, to allow mirror-image imitation by the subject. Test stimuli consisted of 15 meaningless gestures. Performance on each trial was scored by trained coders with established inter-rater reliability (Cohen’s Kappa > 85%) as “correct” (one point) or “incorrect” (zero points) for four aspects of the movement: hand posture, arm posture, amplitude, and timing (Buxbaum et al., 2000). Thus, a maximum of four points could be earned for each gesture (60 total points).

#### Novel Tools Test

The novel tools test assessed the ability to select and use the appropriate novel tool to act upon a target object. This test, a measure of mechanical problem-solving, required participants to select and use novel tools to manipulate novel cylindrical objects (Goldenberg & Hagmann, 1998). Each wooden cylinder had a metal or wooden part to which one of the tools must be fitted (e.g., a “hook” tool fits in the “eye” of a cylinder) to lift the cylinder from a platform. Participants were shown a single cylinder at a time, along with an array of three tools, and asked to select the tool best suited for lifting the cylinder. On each trial, participants received two points for correctly selecting the tool on the first attempt and lost one point each time they made a wrong choice. Once the correct tool was identified, participants were then asked to apply that tool to lift the cylinder from the base. Participants earned two points for successfully lifting the cylinder but lost points for incorrect or failed attempts (scores could not be less than zero for each trial). In our dataset, the selection and application scores were combined to create a Novel Tools Total score (maximum score = 24).

### Attention allocation and control tasks

#### Attention Network Test (ANT)

The attention network test measured different aspects of attention allocation and control, including attention orienting, attention alerting, and conflict resolution (executive score). In the Attention Network Task (ANT; Fan et al., 2002) participants depressed a left or right button on a button box to indicate the orientation of a center arrow presented on a computer monitor while inhibiting a response based on the orientation of flanking arrows. The center arrow direction could be the same (i.e., congruent) or different from the flanking arrows. There was also a neutral condition where the flankers were dashes without any arrowheads. Stimuli appeared either at the bottom or the top of the screen. On some trials, the stimulus could be preceded by a cue orienting participants to the spatial location of the upcoming stimulus (i.e., a star appearing at the top or bottom of the screen). In other trials, the cue alerted individuals that a stimulus was coming but was uninformative about the stimulus location, either presented as a single star appearing at the center of the screen or as stars appearing at both the top and bottom of the screen (i.e., a double cue). Finally, in the remaining trials no cue was presented. Participants were scored on accuracy and reaction time (RT). RT was calculated separately for each of the three congruency conditions (congruent/incongruent/neutral) and four cue location conditions (center/double/spatial cue/no cue), for a total of 12 different mean response times. Within each condition, any RTs that fell three standard deviations above or below the mean were removed, as were any RTs less than 100 ms. Following from Fan et al. (2002), we examined three different scores for analysis. Total accuracy measured a participant’s ability to indicate the orientation of the center arrow and ignore the flanking arrows across all conditions, including both congruent and incongruent trials. The alerting score was computed by subtracting, for correct trials, the RT in the double cue condition from the RT in the no cue condition; this difference was then divided by the average RT of the two conditions. Finally, the orienting score was computed by subtracting, for correct trials, the RT in the spatial cue condition from the RT in the center cue condition, then dividing by the average RT of these two conditions. We were unable to include the executive score (i.e., difference between incongruent and congruent trials, see (Fan et al., 2002) as a predictor as a large proportion of the sample (25%; 13 of 51) performed below chance in the incongruent condition, preventing an accurate estimate of the conflict effect in the reaction times.

#### Digit Span

The digit span is a measure of working memory. This task was taken from the Wechsler Adult Intelligence Scale – III (Weschsler, 1997). Participants had to repeat a gradually increasing sequence of digits spoken at one second intervals by the experimenter, starting with a sequence of two digits. The number of digits in the sequence increased by one after each successful trial. The task terminated when the participant failed two attempts to correctly repeat the sequence of digits. The longest sequence successfully repeated was taken as the digit span measure.

#### Dual-Task Test

The dual-task test is a measure of the ability to resolve interference caused by a cognitive load. In a baseline condition, black dots (1-cm diameter) appeared in random locations on a computer screen at intervals of 0.5, 1.0, 1.5, or 2.0 s. Participants had to press the space bar whenever a dot appeared. In a subsequent dual-task condition, the dot detection task was performed while participants repeated digit strings of length span-minus-one, where the span was taken from the prior Digit Span task. On each trial, the experimenter recited a unique sequence of digits that the participant had to repeat while responding as rapidly as possible to the dot. All trials were considered for analysis regardless of whether the sequence of digits was correctly repeated or not. A dual-task interference score was computed by subtracting the mean baseline RT from the mean dual-task RT (revised from McDowell et al., 1997).

#### Self-Ordered Pointing Test

In this measure of non-verbal working memory, participants were shown twelve black-and-white abstract designs in a 3×4 array on a single sheet of paper. The participants were asked to look over the designs and point to any one of them. The experimenter then turned the page, revealing the same designs in different positions in the array. Participants were asked to point to a different design than any of their previous choices. The test continued in this fashion for twelve trials. Participants received one point each time they chose a design that was different from the previous choices (revised from Petrides and Milner, 1982).

#### Sustained Attention to Response Test (SART)

The SART is a measure of the ability to allocate and sustain attention over an extended time period. In this continuous performance test, 225 single digits were presented serially over a period of 4.3 minutes on a computer screen running E-Prime (Robertson et al., 1997). Participants were asked to respond by pressing a key to all digits except “3,” which appeared 25 times according to a predetermined, quasi-random schedule. The digits varied in size. Each digit was presented for 250 ms, followed by a 900 ms mask. This task was scored as the number of commission errors (i.e., incorrect presses to 3) and the number of omission errors (i.e., lack of response to numbers other than 3). From these two scores we calculated a sensitivity index (d-prime).

### Reasoning Tasks

#### Brixton Test

The Brixton test is a visuospatial sequencing task that measures the ability to detect and follow rules, an aspect of executive functioning. This test (Burgess & Shallice, 1997) employed a testing book consisting of 56 pages (55 trials). On each trial, participants were presented with two rows of five circles each, equally spaced and equal in size. The circles were numbered sequentially based on position. On each page, one circle was colored blue. Participants had to deduce the pattern of movement of the blue circle and predict where the blue circle would appear on the next page. The pattern changed periodically, at which point participants were required to deduce the new pattern and predict the next movement accordingly. The test was scored as the number of trials on which the blue circle’s position was not correctly predicted; better performance is indicated by lower scores.

#### Tower of London

The Tower of London test is a measure of planning ability, an aspect of executive functioning. Participants had to move a stack of balls among three posts to produce a specified configuration, indicated with a colored drawing, within a limited number of moves (Shallice, 1982). There were 12 problems increasing in complexity and number of moves required. Performance was scored as the number of problems solved on the first try and in under 60 seconds.

### Test of Functional Independence

#### Older American Resources and Services – Multidimensional Functional Assessment Questionnaire (OARS-MFAQ)

To examine the relationship of the NAT with impairments in activities of daily living (ADL), participants completed the OARS-MFAQ, a questionnaire measuring functional independence in ADL (Fillenbaum & Smyer, 1981). A subset of seven questions measuring instrumental ADLs (i.e., temporally-extended tasks, such as using the telephone, shopping for groceries, meal preparation, and housework) was administered. Participants were required to select from among three response options: absence of impairment (two points; e.g., “without help, including looking up numbers and dialing a telephone”), some impairment (one point; e.g., “can answer the phone or dial the operator in an emergency, but need a special phone or help in getting the number or dialing”), or inability to perform the task (zero point; e.g., “completely unable to use the telephone”). The total score was calculated from the sum of the seven items, with a maximum score of 14 indicating no ADL impairments.

### Imaging

Of the 51 participants included in this dataset, 45 had a hand-drawn lesion map obtained from a research-quality MRI or CT scan. Participants received their scan at the Hospital of the University of Pennsylvania. Research MRI scans included whole-brain T1-weighted MR images collected on a 3T (Siemens Trio, Erlangen, Germany; repetition time =1620 msec, echo time = 3.87 msec, field of view = 192 × 256 mm, 1 × 1 × 1 mm voxels) or 1.5T (Siemens Sonata, repetition time = 3,000 msec, echo time=3.54msec, field of view= 24 cm, 1.25 × 1.25 × 1.25 mm voxels) scanner, using a Siemens eight-channel head coil. Participants who were contraindicated for MRI underwent whole-brain research CT scans without contrast (60 axial slices, 3–5 mm slice thickness) on a 64-slice Siemens SOMATOM Sensation scanner. For MRI scans, a team member manually segmented lesions to produce a 3-D lesion mask. The binarized lesion drawings were then warped to a 1×1×1 mm common template brain (Montreal Neurological Institute “Colin27”) using a symmetric diffeomorphic registration algorithm (Avants et al., 2008) to translate manual lesion segmentations to standardized space via a two-step process: First, they were registered to an intermediate template comprising healthy brain images acquired from the same scanner at the Hospital of the University of Pennsylvania that was used to collect MRI scans from the patients; then, volumes were mapped from the intermediate template to the “Colin27” template (i.e., Montreal Neurological Institute – MNI – space). A neurologist naive to the behavioral data (Dr. Branch Coslett) inspected all warped lesions to ensure that no errors had occurred. Lesions from research CT or clinical scans were drawn by the same neurologist directly onto the template brain, which had been rotated to match the pitch of the patient’s scan. This method achieved high intra- and interrater reliability in a previous study (Schnur et al., 2009).

### Analyses

#### Least Absolute Shrinkage and Selection Operator (LASSO) Regression

Our dataset consisted of many potential predictors that could explain performance on the NAT. We were interested in assessing which subset of variables meaningfully predicted NAT performance. The challenge of such a large dataset is the presence of collinearities between predictors that cannot be addressed with classic regression models. Thus, we used a LASSO regression approach that has the merit of addressing collinearities. LASSO is a regularized regression technique that applies a penalty term to the residual sum of squares of the regression to achieve a relatively sparse model (Tibshirani, 1996). That is, this penalty term pushes most coefficient (i.e., beta) estimates to zero; any remaining non-zero beta estimates are then assumed to contribute to the model.

Our LASSO regression model was used to test which of 20 variables (Table 4) predicted total NAT performance (i.e., resulted in non-zero model coefficients). Variables included outcome measures from the Behavioral Test Battery, as well as demographic variables including Age and Months post-stroke onset. LASSO regressions were fitted in R using the package *glmnet* (Friedman et al., 2010). Using this package, we first estimated a penalty coefficient (λ) that reflects the amount of shrinkage to apply in the model to minimize mean squared error. Lambda was estimated using a 10-fold cross-validation approach using the *cv.glmnet* function. The estimated minimum λ was used to run the subsequent LASSO regression using the function *glmnet* to estimate the model coefficients. The functions *cv.glmnet* and *glmnet* separately normalize each variable by default, thus accounting for differences in the scale and range of the various battery test scores. Importantly, these functions return non-normalized coefficients that are appropriately scaled for each independent variable (analogous to a conventional regression approach) and do not provide a coefficient uncertainty estimate (e.g., standard error); thus, it is not possible to interpret the magnitude of each coefficient as reflecting its relative importance in the model.

**Table 4:** List of the variables used in the LASSO regression model. In bold are the predictors for which the LASSO regression returned a non-null coefficient.

| *Variable* | *Coefficient estimate* |
| --- | --- |
| **Praxis** |  |
| **Hand Posture Sequence Task** | **-0.0954** |
| **Semantic Gesture Recognition** | **0.0689** |
| Function Knowledge | null |
| Manipulation Knowledge | null |
| Spatial Gesture Recognition | null |
| Gesture to Sight of Object | null |
| Meaningless Imitation | null |
| Novel Tool Use<sup>1</sup> | null |
| **Attention allocation and control** |  |
| **ANT orienting** | **-14.4229** |
| **Dual-Task Interference** | **-0.0004** |
| ANT alerting | null |
| ANT accuracy | null |
| Digit Span | null |
| SOPT | null |
| SART | null |
| **Reasoning** |  |
| Brixton Test | null |
| Tower of London | null |
| **Language** |  |
| **WAB Comprehension** | **0.7245** |
| **Demographics** |  |
| **Age** | **-0.0235** |
| Months Post Stroke onset | null |

<sup>1</sup>The Novel Tool Use task is sometimes referred to as a reasoning task (Goldenberg & Hagmann, 1998), but also relies on praxis system integrity as revealed by correlations in the ability to use novel tools and to use or pantomime familiar tools (Goldenberg & Hagmann, 1998).

#### Prediction quality

Since the LASSO regression method does not provide an estimate of parameter uncertainty, we needed to independently verify that the resulting model (i.e., the non-zero parameters) described the data well. Thus, we performed a leave-one-out cross validation prediction analysis. On each iteration one subject was pulled out of the sample, and model coefficient parameters were estimated using the rest of the sample (i.e., n = 50) using a classic General Linear Model (GLM). This subset model was then used to predict the NAT score of the withheld subject, and the quality of the prediction was compared to the true observed NAT score. This procedure was repeated for all participants in the data set. We applied this procedure twice; once including only the non-zero terms identified from the original LASSO regression, and once including all 20 possible variables. We measured Spearman’s correlation between the actual and predicted NAT scores for each model. To test whether the two Spearman correlations were significantly different (i.e., that the model with non-zero terms was superior to the full model), we applied a permutation approach. We first computed the observed rho difference between each prediction analysis. To test if this difference score was different from a null distribution, we permuted with replacement the NAT scores 10,000 times. For each permutation we calculated the correlation between the predicted NAT scores and the permuted ones under both models, deriving a null distribution of the difference in rho values between the two models. Finally, we estimated the probability that the observed rho difference was null as the number of values from the null distribution that were greater than our true observed rho difference (i.e., the tail probability of observing the actual difference in rho values between the two models). A probability less than 5% (p = 0.05) was considered significant.

#### Correlation between NAT and ADL functional independence questionnaire

To examine the degree to which performance on the NAT relates to an individual’s functional independence in instrumental activities of daily living, we ran a Spearman correlation between the OARS-MFAQ and the NAT.

#### Lesion Symptom-Mapping analyses

To examine lesions associated with poor performance on the NAT, we performed lesion-symptom mapping using Support Vector Regression-Lesion Symptom Mapping (SVR-LSM) using the SVR-LSM Matlab toolbox developed by DeMarco and Turkeltaub (2018). SVR-LSM is a multivariate analysis technique that uses machine learning to determine the association between lesioned voxels and behavior across all voxels and participants simultaneously. Using this approach, we analyzed data from the 45 participants in our dataset that had a hand-drawn lesion map. Only voxels lesioned in at least 10% (i.e., 4) of participants were included in the analysis. Total lesion volume was regressed from both the behavioral scores and the lesion maps. Voxelwise statistical significance was determined using a Monte Carlo style permutation analysis (10,000 iterations) in which the association between behavioral data and lesion map was randomized. A voxelwise threshold of P < 0.005 one-sided was applied to determine chance-level likelihood of a lesion-symptom relationship. A cluster size threshold was subsequently applied to keep only clusters with at least 100 voxels (i.e., 100 mm^3^). The neuroanatomic label of the surviving clusters was checked with the third version of the Automated Anatomical Labelling atlas (AAL3 atlas; Rolls et al., 2019). We report the maximum statistics (Z-value) observed in each cluster, as well as the coordinates of the cluster centroid (Table 5). Finally, using the same approach, we assessed the overlap between the neural underpinnings of the NAT and each of the tasks that emerged as a meaningful predictor of NAT impairments.

**Table 5:** Neural correlates associated with impaired NAT performance. The coordinates for the cluster centroid are in MNI space.

| *Brain area(s)* | *Cluster extent (mm<sup>3</sup>)* | *Peak Z-value* | *Cluster centroid coordinates* |
| --- | --- | --- | --- |
| Inferior Parietal Lobule | 711 | -3.72 | -38; -29; 42 |
| Middle Temporal Gyrus | 339 | -2.78 | -48; -34; -1 |
| Middle Frontal Gyrus | 128 | -3.12 | -32; 35; 22 |

## Results

### NAT performance is predicted by measures of praxis and attention allocation and control

In our retrospective dataset, we first examined the relationship between performance on the Naturalistic Action Test (NAT) and the 20 measures from our background test battery. Using a LASSO regression model, we obtained six non-null predictors (Table 4). These predictors included the number of errors in the Hand Posture Sequencing Task (HPST), the number of items correctly recognized in the Semantic version of the Gesture Recognition Task (Semantic GR), the attention orienting score from the Attention Network Test (ANT orienting), the reaction time difference between the dual-task test and its baseline (hereafter, Dual-Task Interference), the comprehension subtest of the Western Aphasia Battery (WAB Comprehension) and participant age. Importantly, the sign of the coefficient of each predictor matched the expected direction (e.g., positive coefficients for tasks where higher scores indicate better performance). To ensure that these findings were not influenced by the overall stroke severity, we ran a second LASSO regression including NIH stroke scale. This analysis was conducted on the subset of participants for which this score was available (n = 48), as this score was missing for three participants in the database. Crucially, we found that the NIH stroke scale was not a meaningful predictor of NAT performance, while the same six predictors described above survived this new analysis.

To confirm that these six predictors reliably account for NAT performance, we ran a prediction analysis using a leave-one-out cross-validation procedure. Overall, we found that our model predicted actual NAT performance with good accuracy (Spearman correlation between the predicted and actual NAT values, rho = 0.46; p < 0.001 one-sided; Fig. 1A). In comparison, the same cross-validation approach using all 20 predictors led to an inferior prediction of NAT performance (rho = 0.27; p = 0.03 one-sided; Fig. 1B). Using a permutation test, we confirmed that the difference in Spearman’s correlation scores for these two regressions was significantly different (rho difference score = 0.19; p = 0.01 one-sided; Fig. 1C), indicating that the reduced model was a significantly better fit to the data than the full model, potentially due to overfitting by the full model.

**Figure 1:**
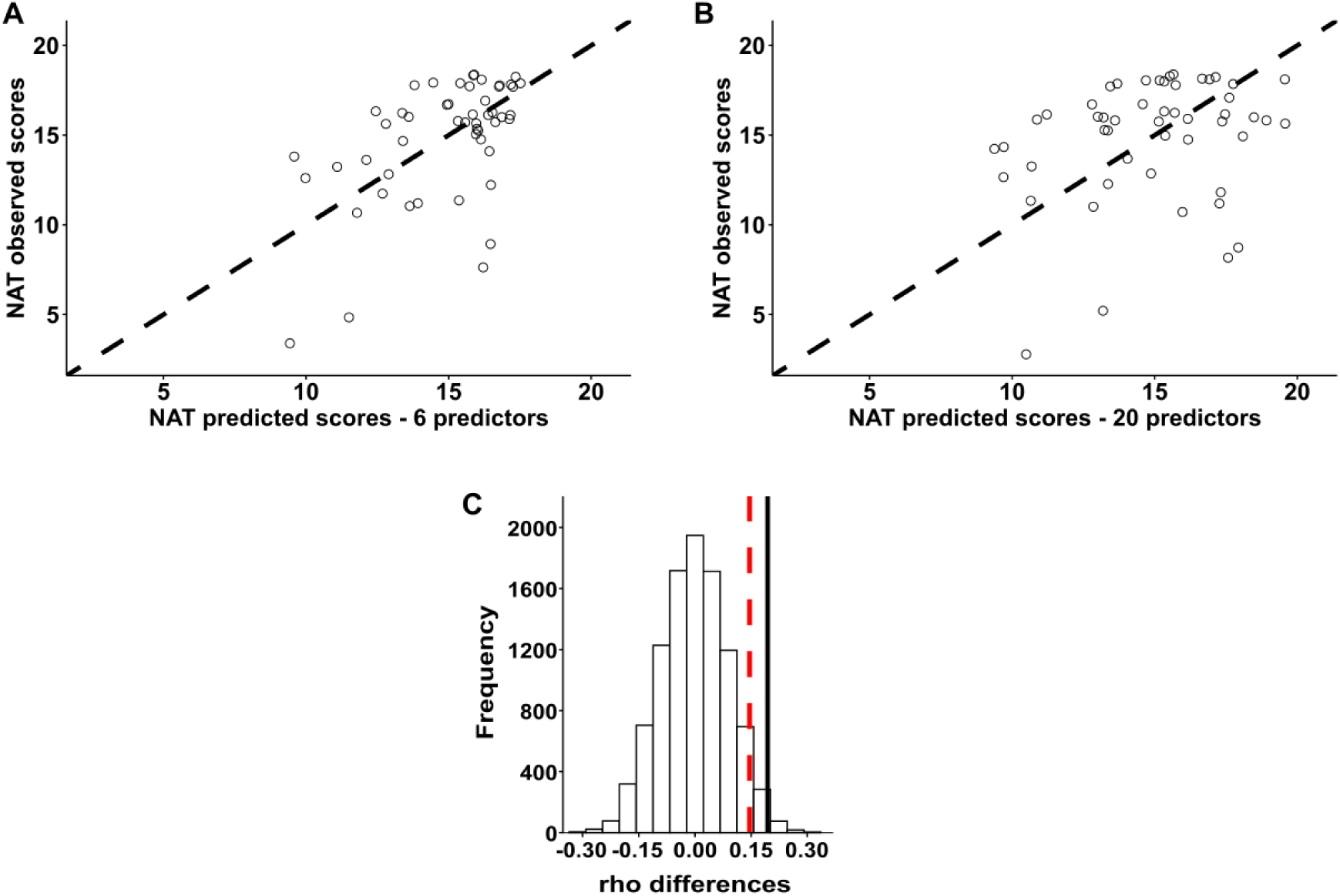
NAT score prediction was better predicted with a classic regression model that included only the non-zero predictors identified by the LASSO regression. Predicted NAT scores were estimated with a classic regression model including A) only the six non-zero predictors from the LASSO regression and, B) all 20 predictors. The black dashed lines depict the unity line. C) The difference between Spearman’s rho scores for each model was calculated and tested for significance compared to a null distribution obtained after 10,000 permutations. The solid black line is the observed rho score difference. The red dashed line reflects an alpha value of 0.05, indicating that the observed rho difference is significantly larger than chance.

### NAT performance relates to impairments in instrumental activities of daily living

Using Spearman’s correlation, we found that NAT performance was significantly correlated with the OARS-MFAQ, a self-report measure of ADL impairments (rho = 0.35; p = 0.01). This positive correlation suggests that participants who performed better on the NAT reported fewer impairments on instrumental activities of daily living.

### Lesions associated with poor NAT performance are associated with praxis and attentional processes

We conducted Support Vector Regression-Lesion Symptom Mapping (SVR-LSM) to examine the relationship between lesion location and performance on the NAT (Fig. 2, Table 5). This analysis indicated that impaired NAT performance was associated with lesions to three brain regions. First, there was an association between impaired NAT performance and left posterior Middle Temporal Gyrus (pMTG). There was also a cluster in the anterior Inferior Parietal Lobule (aIPL), which included voxels in both the Supramarginal Gyrus and the Postcentral Gyrus. Finally, we observed a cluster in the Middle Frontal Gyrus (MFG) extending to the *pars triangularis* of the Inferior Frontal Gyrus (IFG), with 20% of the voxels of that cluster located within the IFG.

**Figure 2:**
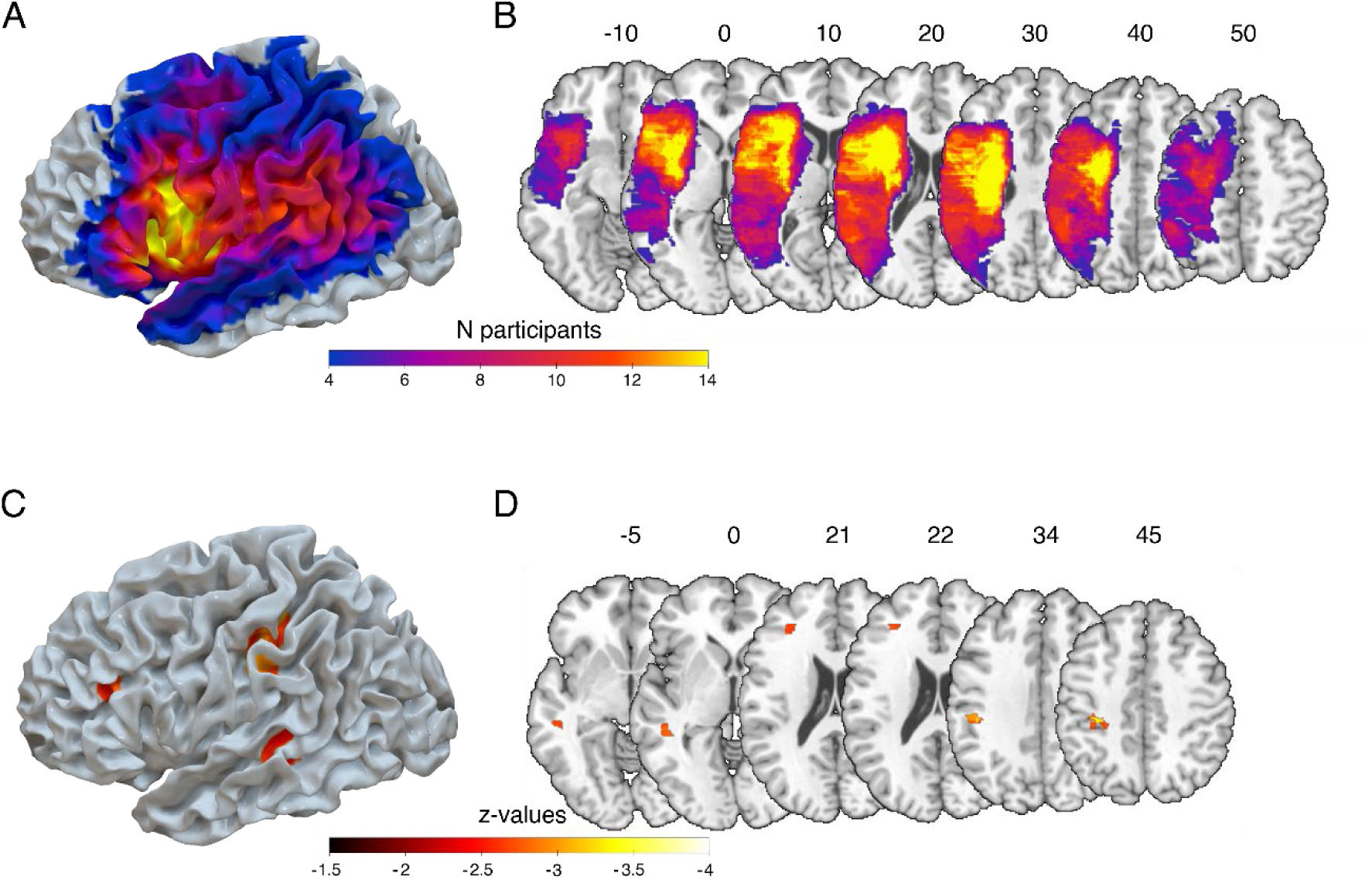
Naturalistic actions impairments are associated with lesions in the frontal, parietal and temporal lobes. Brain surface render (A) and axial-view brain slices (B) are shown for the lesion overlap map depicting voxels lesioned in at least four of the 45 participants; the color reflects the number of individuals for whom that voxel is lesioned (range: 4-14). Slice numbers reflect the corresponding z coordinates in MNI space. Brain surface render (C) and axial brain slices (D) are shown for the thresholded map of areas associated with naturalistic action impairments with significant voxels at p-value < 0.005 one-sided, corresponding to a z-value < −2.57 and a cluster size with at least 100 voxels (i.e., 100 mm^3^). The analysis revealed three main clusters in the pMTG, the aIPL and the MFG.

Finally, we ran a further analysis to explore neuroanatomic overlap between the NAT and the tasks that predicted NAT impairments (Fig. S1). We found that impairments in both the semantic gesture recognition and the ANT orienting score were associated with temporal lobe lesions overlapping with voxels associated with NAT performance in the pMTG (overlap extent: 126 mm^3^ for semantic gesture recognition; 339 mm^3^ for ANT orienting). Dual-Task Interference impairments were associated with lesion in the MFG, overlapping with the MFG cluster found for NAT impairments (overlap extent: 19 mm^3^). In contrast, no overlap was found between lesions associated with errors in the hand-posture sequencing task and the NAT. Finally, both language comprehension (i.e., WAB) and NAT impairments overlapped in the pMTG (overlap extent: 339 mm^3^) and the aIPL (overlap extent: 202 mm^3^).

## Discussion

Based on a retrospective analysis of previously-collected data, we examined the predictors of naturalistic action impairments in a LCVA population using a regularized regression approach (LASSO regression) that accounted for collinearities in the dataset. As expected, we found that naturalistic action impairments in people with LCVA were predicted by praxis deficits. We also found that naturalistic action impairments in this population were predicted by deficits in attentional allocation and control. Finally, language comprehension and age were also predictive of NAT performance.

Praxis is an important contributor to naturalistic action performance. Specifically, we found naturalistic action performance was driven by the ability to recognize tool-related gestures – a measure of praxis knowledge, and the ability to properly produce a series of object-directed hand postures – a measure of sequence production. Regarding tool-use gestures, we specifically observed that the inability to distinguish between semantically-related tool-use gestures (e.g., use of hammer versus saw) was predictive of naturalistic action deficits. Loss of knowledge of the actions associated with manipulable objects has historically been viewed as a core component of so-called “ideational apraxia” (e.g., De Renzi and Lucchelli, 1988; Ochipa et al., 1989). In naturalistic tasks, appropriate objects must be selected in a timely manner from arrays that often contain task-irrelevant but semantically-related objects (e.g., various kitchen utensils in a breakfast preparation task). We speculate that degradation of tool action knowledge makes such selection difficult, contributing to poor task performance. In contrast, we found that the ability to use novel tools was not predictive of naturalistic action performance. These findings are strongly consistent with the results of a prior study showing that familiar tool use, unlike the selection and application of novel tools, correlated with performance of a naturalistic action test based upon the NAT (Buchmann & Randerath, 2017). We also found that the ability to produce a sequence of object-related hand postures plays a role in naturalistic action impairment. Previous studies have shown that individuals with LCVA are particularly impaired at sequencing postures compared to neurotypicals and individuals with RCVA (Haaland & Harrington, 1994; Harrington & Haaland, 1992, 1991). The Hand Posture Sequencing Task requires translation from a sequence of semi-abstract visual symbols to a sequence of hand postures. Poor performance may reflect slow retrieval of the individual hand-posture elements within each sequence and/or a deficit in representing the order of the elements within sequences. We speculate that hand posture sequencing is relevant to naturalistic action by virtue of the fact that in naturalistic action tasks, visual information (i.e., an object in the array) is used to inform the appropriate selection of a hand posture for use of that object. Moreover, naturalistic tasks, like the hand posture sequencing task, contain numerous objects for which the correct hand posture (e.g., cutting butter, spreading butter, and spreading jam with a knife) must be produced sequentially (Botvinick & Plaut, 2004; R. P. Cooper et al., 2005; R. Cooper & Shallice, 2000). Thus, it is possible that naturalistic action suffers from a reduced ability to produce actions in the correct order, over and above any difficulties with retrieval of the individual elements in the sequence. Further investigation is needed to disentangle the precise role of hand posture sequencing deficits in naturalistic actions.

In addition to praxis deficits, we found that attentional allocation and control deficits were also predictive of naturalistic action impairments. Specifically, we observed a relationship with the ability to select information from visual input (orienting) and the ability to divide attention between two simultaneous tasks (dual-task decrements). The orienting measure suggests that a reduced ability to respond to exogenous cues to orient spatial attention contributed to naturalistic action errors. A previous report showed that attention orienting of the Attention Network Test was disrupted in people with LCVA compared to neurotypicals (LaCroix et al., 2020). Attention orienting is typically conceptualized as occurring in response to suddenly-appearing stimuli (e.g., Posner, 1980). While sudden changes of state do occasionally occur in the NAT (e.g., the toast pops up when toasting is complete), in the context of naturalistic tasks we speculate that reductions in orienting attention more broadly impact the degree to which any object in the array provides a “bottom-up” signal of the need to shift attention to its location.

We also found that naturalistic action impairments were related to attention control; specifically, the ability to respond to stimuli despite interference by a secondary task. As noted in the Introduction, a previous study showed that neurotypical individuals made more errors when required to perform a naturalistic action in parallel with another attention-demanding task (Giovannetti et al., 2007). In the naturalistic action tasks performed in the present study, there was no explicit secondary cognitive task. However, naturalistic actions inherently require the division of attention across multiple simultaneously-occurring events. For instance, when preparing breakfast, one can slice butter or open a jam container while simultaneously attending to the status of bread in the toaster. Consistent with prior evidence of dual-task impairments in stroke (Feld & Plummer, 2021; Mullick et al., 2021), deficits in the ability to divide attention across the different subparts of naturalistic action tasks may be impaired in the LCVA population.

It is reasonable to assume that task novelty may play a role in the degree to which attentional resources are taxed. A previous study suggested that naturalistic actions with which most people have little experience (e.g., fixing a video recorder) are correlated with the ability to solve mechanical problems (Hartmann et al., 2005). In contrast, more familiar naturalistic actions (e.g., preparing a cup of coffee) are correlated with praxis and semantic knowledge, in keeping with the results of the present study. However, attention was not measured in that prior study. Future studies may be informative in assessing whether task familiarity mediates dependence on attentional processes.

We additionally found that language comprehension is predictive of naturalistic action performance. It is worth noting that the NAT is designed to limit verbal instructions by showing people visual pictures of the task goal (e.g., a picture of toast with butter and jam) and also includes a phase where the experimenter checks whether the instructions are understood. Despite these methodological considerations, we cannot fully reject the possibility that some language comprehension deficits affected task performance, as both the NAT and parts of the WAB involve remembering and following complex multi-step instructions. This account is consistent with a previous study in individuals with LCVA that found a correlation between measures of language comprehension and performance in a coffee preparation task (Hartmann et al., 2005; see also Stoll et al., 2022). Finally, we found that (older) age was also predictive of greater naturalistic action impairments. In line with our findings, a previous study in neurotypicals found that the number of errors in naturalistic action is greater for older participants compared to younger ones (Giovannetti et al., 2019). This result is perhaps unsurprising given that aging affects overall brain health, which in turn affects “general” cognitive functions (Brayne et al., 1995).

In addition to examining the predictors of NAT performance, we also assessed whether the Naturalistic Action Test is a valid measure of participants’ assessment of their abilities in temporally-extended activities of daily living. We observed a moderate correlation between the NAT and self-reported responses on the OARS-MFAQ. On that basis, we suggest that the results of this study have relevance not just for laboratory-based tests of naturalistic actions, but for real-world activities more broadly.

Lastly, we conducted a lesion-symptom mapping analysis to explore which brain regions may be involved in naturalistic action deficits. Our analysis revealed that lesions to portions of the praxis (tool-use) network including the pMTG, the aIPL and the MFG are associated with naturalistic action impairments. The contribution of these areas is consistent with our behavioral data showing that praxis deficits predict naturalistic action impairments, and with previous neuroimaging studies in which neurotypicals observed naturalistic actions (see Bienkiewicz et al., 2014 for a meta-analysis). The left pMTG plays a role in action observation and recognition (Papeo et al., 2019; Wurm & Lingnau, 2015) and subserves representations for tools and hands; lesions to the left pMTG contribute to impaired gesture knowledge (Kalénine et al., 2010; Tarhan et al., 2015; see Fig. S1 for Semantic Gesture Recognition). Based on our finding that impaired semantic gesture recognition contributed to poorer naturalistic action performance, we suggest that lesions to the pMTG contribute to degraded tool-use representations that impact naturalistic action. Lesions to the aIPL, in contrast, are associated with errors in the production of hand postures in tool-use pantomime (Hoeren et al., 2014; Tarhan et al., 2015). While in our study tool-use pantomime (i.e. Gesture-to-Sight of Object task) was not a contributing predictor to naturalistic action impairments, we may speculate that lesions to the aIPL contribute to an inability to select hand postures appropriate to objects in the naturalistic array.

The MFG is known to be involved in the execution of tool-related actions (Brandi et al., 2014; Chen et al., 2023; Gallivan et al., 2013), although the specific cognitive process supported by this area remains unclear. As demonstrated by our overlap analysis (Fig. S1 for Dual-Task Interference), the MFG is also involved in dual-task performance (Worringer et al., 2019). Thus, this area may contribute to both attention and praxis computations that are critical for naturalistic action.

Note that the use of lesion-symptom mapping techniques is limited by the set of lesion locations (i.e., the lesion map overlap) within a sample, which in our case tends to be sparse for more posterior areas such the occipito-temporal cortex involved in praxis (Brandi et al., 2014; Peeters et al., 2009) and for prefrontal areas involved in attentional control (Dove et al., 2000; MacDonald et al., 2000). Functional connectivity analyses in people with LCVA or neurotypicals may be useful in further clarifying the brain networks involved in the performance of naturalistic action.

Despite the strength and the novelty of the multivariate LASSO regression approach employed, the current study has some limitations. In particular, the analysis of a retrospective dataset limited our access to fine-grained details about administration and data pre-processing for some tasks, as well as trial-level data that might allow for more sophisticated statistical modeling approaches. Future research can address these limitations by taking advantage of recent progress in the development of large publicly-available datasets that can be analyzed with contemporary statistical approaches.

Taken together, findings from this study provide evidence that naturalistic action impairments in people with LCVA are related to praxis deficits, as well as deficits in attention allocation and control, language comprehension, and age. These data contradict the traditional view in the praxis literature that naturalistic action tasks are simple extensions of single object pantomime and use tasks (De Renzi & Lucchelli, 1988; Poeck, 1983), and help to explain why naturalistic action impairments are also observed in people with right hemisphere stroke, traumatic brain injury, and dementia. The current findings point to several cognitive processes that play a key role in the successful production of naturalistic actions, including knowledge of object manipulation, the ability to produce object-appropriate hand postures in response to environmental input, the allocation of attention in response to visual cues, and the ability to resist cognitive interference. Importantly, our study confirms the role of attentional factors in performing naturalistic actions. Future studies are needed to compare the degree to which the performance of people with left hemisphere and right hemisphere stroke (as well as other disorders such as traumatic brain injury and dementia) are influenced by such attentional factors.

## Acknowledgements

Data were collected with funding from the National Institute of Disability and Rehabilitation Research to LJB (H133G030169). We thank Myrna Schwartz for her invaluable input, Susan Lipsett for help with data collection, Branch Coslett for assistance with brain lesion segmentation, and Andrew DeMarco for help with the SVR-LSM analyses. This work is supported by NIH grant R01 NS115862 awarded to ALW.

## References

Anderson, B. A. (2021). An Adaptive View of Attentional Control. American Psychologist, 76(9), 1410–1422. 10.1037/amp0000917

Avants, B. B., Epstein, C. L., Grossman, M., & Gee, J. C. (2008). Symmetric Diffeomorphic Image Registration with Cross-Correlation: Evaluating Automated Labeling of Elderly and Neurodegenerative Brain. Medical Image Analysis, 12(1), 26–41. www.itk.org

Balouch, S., & Rusted, J. M. (2014). Error-monitoring in an everyday task in people with Alzheimer-type dementia: Observations over five years of performance decline. Journal of Clinical and Experimental Neuropsychology, 36(7), 773–786. 10.1080/13803395.2014.943697

Bickerton, W.-L., Humphreys, G. W., & Riddoch, M. J. (2007). The case of the unfamiliar implement: Schema-based over-riding of semantic knowledge from objects in everyday action. Journal of the International Neuropsychological Society, 13, 1035–1047. 10.1017/S1355617707071585

Bickerton, W.-L., Riddoch, M. J., Samson, D., Balani, A. B., Mistry, B., & Humphreys, G. W. (2012). Systematic assessment of apraxia and functional predictions from the Birmingham Cognitive Screen. Cognitive Neurology, 83, 513–521. 10.1136/jnnp-2011-300968

Bienkiewicz, M. M. N., Brandi, M.-L., Goldenberg, G., Hughes, C. M. L., Hermsdörfer, J., & Binkofski, F. (2014). The tool in the brain: apraxia in ADL. Behavioral and neurological correlates of apraxia in daily living. Frontiers in Psychology, 5(353), 1–13. 10.3389/fpsyg.2014.00353

Bienkiewicz, M. M. N., Brandi, M.-L., Hughes, C., Voitl, A., & Hermsdörfer, J. (2015). The complexity of the relationship between neuropsychological deficits and impairment in everyday tasks after stroke. Brain and Behavior, 5(10), 1–14. 10.1002/brb3.371

Binkofski, F., & Buxbaum, L. J. (2013). Two action systems in the human brain. Brain and Language, 127(2), 222–229. 10.1016/j.bandl.2012.07.007

Borod, J. C., Carper, M., Naeser, M., & Goodglass, H. (1985). Left-handed and right-handed aphasics with left hemisphere lesions compared on nonverbal performance measures. Cortex, 21, 81–90.

Boronat, C. B., Buxbaum, L. J., Coslett, B. H., Tang, K., Saffran, E. M., Kimberg, D. Y., & Detre, J. A. (2005). Distinctions between manipulation and function knowledge of objects: evidence from functional magnetic resonance imaging. Cognitive Brain Research, 23, 361–373. 10.1016/j.cogbrainres.2004.11.001

Boshra, R., & Kastner, S. (2022). Attention control in the primate brain This review comes from a themed issue on Systems Neuroscience. Current Opinion in Neurobiology, 2022, 102605. 10.1016/j.conb.2022.102605

Botvinick, M., & Plaut, D. C. (2004). Doing Without Schema Hierarchies : A Recurrent Connectionist Approach to Normal and Impaired Routine Sequential Action. Psychological Review, 111(2), 395–429. 10.1037/0033-295X.111.2.395

Brandi, M.-L., Wohlschläger, A., Sorg, C., & Hermsdörfer, J. (2014). The Neural Correlates of Planning and Executing Actual Tool Use. The Journal of Neuroscience, 34(39), 13183–13194. 10.1523/JNEUROSCI.0597-14.2014

Brayne, C., Gill, C., Paykel, E. S., Huppert, F., & O’connor, D. W. (1995). Cognitive decline in an elderly population-a two wave study of change. Psychological Medicine, 25, 673–683.

Brott, T., Adams, H. P., Olinger, C. P., Marle, J. R., Barsan, W. G., Biller, J., Spilker, J., Holleran, R., Eberle, R., Hertzberg, V., Rorick, M., Moomaw, C. J., & Walker, M. (1989). Measurements of acute cerebral infarction: A clinical examination scale. Stroke, 20(7), 864–870. 10.1161/01.STR.20.7.864

Buchmann, I., & Randerath, J. (2017). Selection and application of familiar and novel tools in patients with left and right hemispheric stroke: Psychometrics and normative data. Cortex, 94, 49–62. 10.1016/j.cortex.2017.06.001

Burgess, P. W., & Shallice, T. (1997). The relationship between prospective and retrospective memory: Neuropsychological evidence. In M. Press (Ed.), Cognitive models of memory (M.A. Conwa, pp. 247–272).

Burgoyne, A. P., & Engle, R. W. (2020). Attention Control: A Cornerstone of Higher-Order Cognition. Current Directions in Psychological Science, 29(6), 624–630. 10.1177/0963721420969371

Buxbaum, L. J., Giovannetti, T., & Libon, D. (2000). The Role of the Dynamic Body Schema in Praxis: Evidence from Primary Progressive Apraxia. Brain and Cognition, 44, 166–191. 10.1006/brcg.2000.1227

Buxbaum, L. J., & Kalénine, S. (2010). Action knowledge, visuomotor activation, and embodiment in the two action systems. Annals of the New York Academy of Sciences, 1191(1), 201–218. 10.1111/J.1749-6632.2010.05447.X

Buxbaum, L. J., & Kalénine, S. (2022). Apraxia : A disorder at the cognitive-motor interface. In M. S. Gazzaniga & G. R. Mangun (Eds.), The Cognitive Neurosciences (5th ed.).

Buxbaum, L. J., Kyle, K. M., & Menon, R. (2005). On beyond mirror neurons: Internal representations subserving imitation and recognition of skilled object-related actions in humans. 10.1016/j.cogbrainres.2005.05.014

Buxbaum, L. J., Schwartz, M. F., & Montgomery, M. W. (1998). Ideational Apraxia and Naturalistic Action. Cognitive Neuropsychology, 15(6–8), 617–643. 10.1080/026432998381032

Chen, J., Paciocco, J. U., Deng, Z., & Culham, J. C. (2023). Human Neuroimaging Reveals Differences in Activation and Connectivity between Real and Pantomimed Tool Use. Journal of Neuroscience, 43(46), 7853–7867. 10.1523/JNEUROSCI.0068-23.2023

Cooper, R. P., Schwartz, M. F., Yule, P., & Shallice, T. (2005). The simulation of action disorganisation in complex activities of daily living. Cognitive Neuropsychology, 22(8), 959–1004. 10.1080/02643290442000419

Cooper, R., & Shallice, T. (2000). Contention scheduling and the control of routine activities. Cognitive Neuropsychology, 17(4), 297–338. 10.1080/026432900380427

De Renzi, E., & Lucchelli, F. (1988). Ideational Apraxia. Brain, 111, 1173–1185.

DeMarco, A. T., & Turkeltaub, P. E. (2018). A multivariate lesion symptom mapping toolbox and examination of lesion-volume biases and correction methods in lesion-symptom mapping. Human Brain Mapping, 39(11), 4169–4182. 10.1002/hbm.24289

Dove, A., Pollmann, S., Schubert, T., Wiggins, C. J., & Yves Von Cramon, D. (2000). Prefrontal cortex activation in task switching: an event-related fMRI study. Cognitive Brain Research, 9, 103–109. www.elsevier.comrlocaterbres

Fan, J., McCandliss, B. D., Fossella, J., Flombaum, J. I., & Posner, M. I. (2005). The activation of attentional networks. NeuroImage, 26(2), 471–479. 10.1016/j.neuroimage.2005.02.004

Fan, J., McCandliss, B. D., Sommer, T., Raz, A., & Posner, M. I. (2002). Testing the efficiency and independence of attentional networks. Journal of Cognitive Neuroscience, 14(3), 340–347. 10.1162/089892902317361886

Feld, J. A., & Plummer, P. (2021). Patterns of cognitive-motor dual-task interference post stroke: An observational inpatient study at hospital discharge. European Journal of Physical and Rehabilitation Medicine, 57(3), 327–336. 10.23736/S1973-9087.20.06273-5

Fillenbaum, G. G., & Smyer, M. A. (1981). The development, validity, and reliability of the OARS multidimensional functional assessment questionnaire. Journals of Gerontology, 36(4), 428–434. 10.1093/geronj/36.4.428

Friedman, J., Hastie, T., & Tibshirani, R. (2010). Regularization Paths for Generalized Linear Models via Coordinate Descent. JSS Journal of Statistical Software, 33. http://www.jstatsoft.org/

Gallivan, J. P., Adam McLean, D., Valyear, K. F., & Culham, J. C. (2013). Decoding the neural mechanisms of human tool use. ELife, 2013(2), 1–29. 10.7554/eLife.00425

Giovannetti, T., Bettcher, B. M., Brennan, L., Libon, D. J., Kessler, R. K., & Duey, K. (2008). Coffee With Jelly or Unbuttered Toast : Commissions and Omissions Are Dissociable Aspects of Everyday Action Impairment in Alzheimer ’ s Disease. Neuropsychology, 22(2), 235–245. 10.1037/0894-4105.22.2.235

Giovannetti, T., Britnell, P., Brennan, L., Siderowf, A., Grossman, M., Libon, D. J., Bettcher, B. M., Rouzard, F., Eppig, J., & Gregory A., S. (2012). Everyday Action Impairment in Parkinson ’ s Disease Dementia. Journal of the International Neuropsychology Society, 18(5), 787–798. 10.1017/S135561771200046X.Everyday

Giovannetti, T., Libon, D. J., Buxbaum, L. J., & Schwartz, M. F. (2002). Naturalistic action impairments in dementia. Neuropsychologia, 40, 1220–1232.

Giovannetti, T., Schwartz, M. F., & Buxbaum, L. J. (2007). The Coffee Challenge: A new method for the study of everyday action errors. Journal of Clinical and Experimental Neuropsychology, 29(7), 690–705. 10.1080/13803390600932286

Giovannetti, T., Yamaguchi, T., Roll, E., Harada, T., Rycroft, S. S., Divers, R., Hulswit, J., Tan, C. C., Matchanova, A., Ham, L., Hackett, K., & Mis, R. (2019). The Virtual Kitchen Challenge: preliminary data from a novel virtual reality test of mild difficulties in everyday functioning. Aging, Neuropsychology, and Cognition, 26(6), 823–841. 10.1080/13825585.2018.1536774

Goldenberg, G. (2013). Apraxia in left-handers. Brain, 136, 2592–2601. 10.1093/brain/awt181

Goldenberg, G., & Hagmann, S. (1997). The meaning of meaningless gestures: A study of visuo-imitative apraxia. Neuropsychologia, 35(3), 333–341.

Goldenberg, G., & Hagmann, S. (1998). Tool use and mechanical problem solving in apraxia. Neuropsychologia, 36(7), 581–589.

Gonzalez Rothi, L. J., Heilman, K. M., & Watson, R. T. (1985). Pantomime comprehension and ideomotor apraxia. Journal of Neurology Neurosurgery and Psychiatry, 48(3), 207–210. 10.1136/jnnp.48.3.207

Haaland, K. Y., & Harrington, D. L. (1994). Limb-Sequencing Deficits after Left but not Right Hemisphere Damage. Brain and Cognition, 24, 104–122.

Harrington, D. L., & Haaland, K. (1992). Motor sequencing with left hemisphere damage are some cognitive deficits specific to limb apraxia. Brain, 115, 857–874.

Harrington, D. L., & Haaland, K. Y. (1991). Hemispheric specialization for motor sequencing: abnormalities in levels of programming. Neuropsychologia, 2(147– 163), 147–163. 10.1016/0028-3932(91)90017-3

Hartmann, K., Goldenberg, G., Daumüller, M., & Hermsdörfer, J. (2005). It takes the whole brain to make a cup of coffee: the neuropsychology of naturalistic actions involving technical devices. Neuropsychologia, 43, 625–637. 10.1016/j.neuropsychologia.2004.07.015

Heilman, K. M., Rothi, L. J., & Valenstein, E. (1982). Two forms of ideomotor apraxia. Neurology, 32(4), 342–446. 10.1212/wnl.32.4.342

Hoeren, M., Kü, D., Bormann, T., Beume, L., Ludwig, V. M., Vry, M.-S., Mader, I., Rijntjes, M., Kaller, C. P., & Weiller, C. (2014). Neural bases of imitation and pantomime in acute stroke patients: distinct streams for praxis. Brain, 137, 2796– 2810. 10.1093/brain/awu203

Humphreys, G. W., & Riddoch, J. M. (2000). One more cup of coffee for the road: Object-action assemblies, response blocking and response capture after frontal lobe damage. In Experimental Brain Research (Vol. 133, Issue 1, pp. 81–93). Springer Verlag. 10.1007/s002210000403

Jarry, C., Osiurak, F., Baumard, J., Lesourd, M., Coiffard, C., Lucas, C., Merck, C., Etcharry-Bouyx, F., Chauviré, V., Belliard, S., Moreaud, O., Croisile, B., & Le Gall, D. (2021). Daily life activities in patients with Alzheimer’s disease or semantic dementia: Multitasking assessment. Neuropsychologia, 150. 10.1016/j.neuropsychologia.2020.107714

Johnson-Frey, S. H. (2004). The neural bases of complex tool use in humans. Trends in Cognitive Sciences, 8(2), 71–78. 10.1016/j.tics.2003.12.002

Kalénine, S., Buxbaum, L. J., & Coslett, B. (2010). Critical brain regions for action recognition: lesion symptom mapping in left hemisphere stroke. Brain, 133, 3269– 3280. 10.1093/brain/awq210

Kertesz, A., & Hooper, P. (1982). Praxis and language: the extent and variety of apraxia in aphasia. Neuropsychologia, 20(3), 275–286. 10.1016/0028-3932(82)90102-6

Kroliczak, G., Buchwald, M., Kleka, P., Klichowski, M., Potok, W., Nowik, A. M., Randerath, J., & Piper, B. J. (2021). Manual praxis and language-production networks, and their links to handedness. Cortex, 140, 110–127. 10.1016/j.cortex.2021.03.022

LaCroix, A. N., Tully, M., & Rogalsky, C. (2020). Assessment of alerting, orienting, and executive control in persons with aphasia using the Attention Network Test. Aphasiology, 00(00), 1–16. 10.1080/02687038.2020.1795077

Lewis, J. W. (2006). Cortical networks related to human use of tools. Neuroscientist, 12(3), 211–231. 10.1177/1073858406288327

MacDonald, A. W., Cohen, J. D., Andrew Stenger, V., & Carter, C. S. (2000). Dissociating the role of the dorsolateral prefrontal and anterior cingulate cortex in cognitive control. Science, 288(5472), 1835–1838. 10.1126/SCIENCE.288.5472.1835

Manohar, S., Bonnelle, V., & Husain, M. (2014). Neurological disorders of attention. In C. Nobre & S. Kastner (Eds.), The Oxford handbook of attention (University, pp. 1028–1061).

McDowell, S., Whyte, J., & D’Esposito, M. (1997). Working memory impairments in traumatic brain injury: evidence from a dual-task paradigm. 35(10), 1341–1353.

Miller, E. K., & Cohen, J. D. (2001). An Integrative Theory of Prefrontal Cortex Function. Annual Review of Neuroscience, 24, 167–202.

Mullick, A. A., Baniña, M. C., Tomita, Y., Fung, J., & Levin, M. F. (2021). Obstacle Avoidance and Dual-Tasking During Reaching While Standing in Patients With Mild Chronic Stroke. Neurorehabilitation and Neural Repair, 35(10), 915–928. 10.1177/15459683211023190

Ochipa, C., Gonzalez Rothi, L. J., & Heilman, K. M. (1989). Ideational apraxia: a deficit in tool selection and use. Annals of Neurology, 25(2), 190–193. 10.1002/ana.410250214

Orban, G. A., & Caruana, F. (2014). The neural basis of human tool use. Frontiers in Psychology, 5(310), 1–12. 10.3389/fpsyg.2014.00310

Papeo, L., Agostini, B., & Lingnau, A. (2019). The Large-Scale Organization of Gestures and Words in the Middle Temporal Gyrus. Journal of Neuroscience, 39(30), 5966– 5974. 10.1523/JNEUROSCI.2668-18.2019

Peeters, R. R., Simone, L., Nelissen, K., Fabbri-Destro, M., Vanduffel, W., Rizzolatti, G., & Orban, G. A. (2009). The Representation of Tool Use in Humans and Monkeys: Common and Uniquely Human Features. Journal of Neuroscience, 29(37), 11523– 11539. 10.1523/JNEUROSCI.2040-09.2009

Petersen, S. E., & Posner, M. I. (2012). The Attention System of the Human Brain: 20 Years After. Annual Review of Neurpscience, 35, 73–89. 10.1146/annurev-neuro-062111-150525

Petrides, M., & Milner, B. (1982). Deficits on subject-ordered tasks after frontal- and temporal-lobe lesions in man. Neuropsychologia, 20(3), 249–262. 10.1016/0028-3932(82)90100-2

Poeck, K. (1983). Ideational apraxia. Journal of Neurology, 230, 1–5.

Poole, J. L., Sadek, J., & Haaland, K. Y. (2011). Meal Preparation Abilities After Left or Right Hemisphere Stroke. Archives of Physical Medicine and Rehabilitation, 92(4), 590–596. 10.1016/j.apmr.2010.11.021

Posner, M. I. (1980). Orienting of attention. The Quarterly Journal of Experimental Psychology, 32(1), 3–25. 10.1080/00335558008248231

Reason, J. (1984). Lapses of Attention in Everyday Life. In R. Parasuraman & D. Davies (Eds.), Varieties of attention (pp. 515–549). Academic Press.

Robertson, I. H., Manly, T., Andrade, J., Baddeley, B. T., & Yiend, J. (1997). “Oops!”: Performance correlates of everyday attentional failures in traumatic brain injured and normal subjects. Neuropsychologia, 35(6), 747–758.

Roll, E. E., Giovannetti, T., Libon, D. J., & Eppig, J. (2017). Everyday task knowledge and everyday function in dementia. Journal of Neuropsychology, 13(1), 1–25. 10.1111/jnp.12135

Rolls, E. T., Huang, C.-C., Lin, C.-P., Feng, J., & Joliot, M. (2019). Automated anatomical labelling atlas 3. NeuroImage, 116189, 1–5. 10.1016/j.neuroimage.2019.116189

Rumiati, R. I. (2005). Right, left or both? Brain hemispheres and apraxia of naturalistic actions. Trends in Cognitive Sciences, 9(4), 164–167. 10.1016/j.tics.2005.02.001

Schnur, T. T., Schwartz, M. F., Kimberg, D. Y., Hirshorn, E., Branch Coslett, H., & Thompson-Schill, S. L. (2009). Localizing interference during naming: Convergent neuroimaging and neuropsychological evidence for the function of Broca’s area. In PNAS (Vol. 106, Issue 1).

Schwartz, M. F., Buxbaum, L. J., Montgomery, M. W., Fitzpatrick-Desalme, E., Hart, T., Ferraro, M., Lee, S. S., & Coslett, B. H. (1999). Naturalistic action production following right hemisphere stroke. Neuropsychologia, 37, 51–66.

Schwartz, M. F., Lee, S. S., Coslett, H. B., Montgomery, M. W., Buxbaum, L. J., Carew, T. G., Ferraro, M., Fitzpatrick-DeSalme, E., Hart, T., & Mayer, N. (1998). Naturalistic action impairment in closed head injury. Neuropsychology, 12(1), 13–28. 10.1037/0894-4105.12.1.13

Schwartz, M. F., Segal, M., Veramonti, T., Ferraro, M., & Buxbaum, L. J. (2002). The Naturalistic Action Test: A standardised assessment for everyday action impairment. Neuropsychological Rehabilitation, 12(4), 311–339. 10.1080/09602010244000084

Shallice, T. (1982). Specific impairments of planning. In Trans. R. Soc. Lond. B (Vol. 298).

Stoll, S. E. M., Finkel, L., Buchmann, I., Hassa, T., Spiteri, S., Liepert, J., & Randerath, J. (2022). 100 years after Liepmann–Lesion correlates of diminished selection and application of familiar versus novel tools. Cortex, 146, 1–23. 10.1016/j.cortex.2021.10.002

Tarhan, L. Y., Watson, C. E., & Buxbaum, L. J. (2015). Shared and Distinct Neuroanatomic Regions Critical for Tool-related Action Production and Recognition: Evidence from 131 Left-hemisphere Stroke Patients. Journal of Cognitive Neuroscience, 27(12), 2491–2511. 10.1162/jocn_a_00876

Tibshirani, R. (1996). Regression Shrinkage and Selection via the Lasso. Journal of the Royal Statistical Society. Series B (Methodological), 58(1), 267–288.

Weschsler, D. (1997). Wechsler Adult Intelligence Scale - III (Psychological Corporation, Ed.; 3rd ed.).

Worringer, B., Langner, R., Koch, I., Simon, ·, Eickhoff, B., Eickhoff, C. R., & Binkofski, F. C. (2019). Common and distinct neural correlates of dual-tasking and task-switching: a meta-analytic review and a neuro-cognitive processing model of human multitasking. Brain Structure and Function, 224, 1845–1869. 10.1007/s00429-019-01870-4

Wurm, M., & Lingnau, A. (2015). Decoding Actions at Different Levels of Abstraction. Journal of Neuroscience, 35(20), 7727–7735. 10.1523/JNEUROSCI.0188-15

