## Supplementary Materials for "Cognitive neuropsychological and neuroanatomic predictors of naturalistic action performance in left hemisphere stroke: a retrospective analysis"

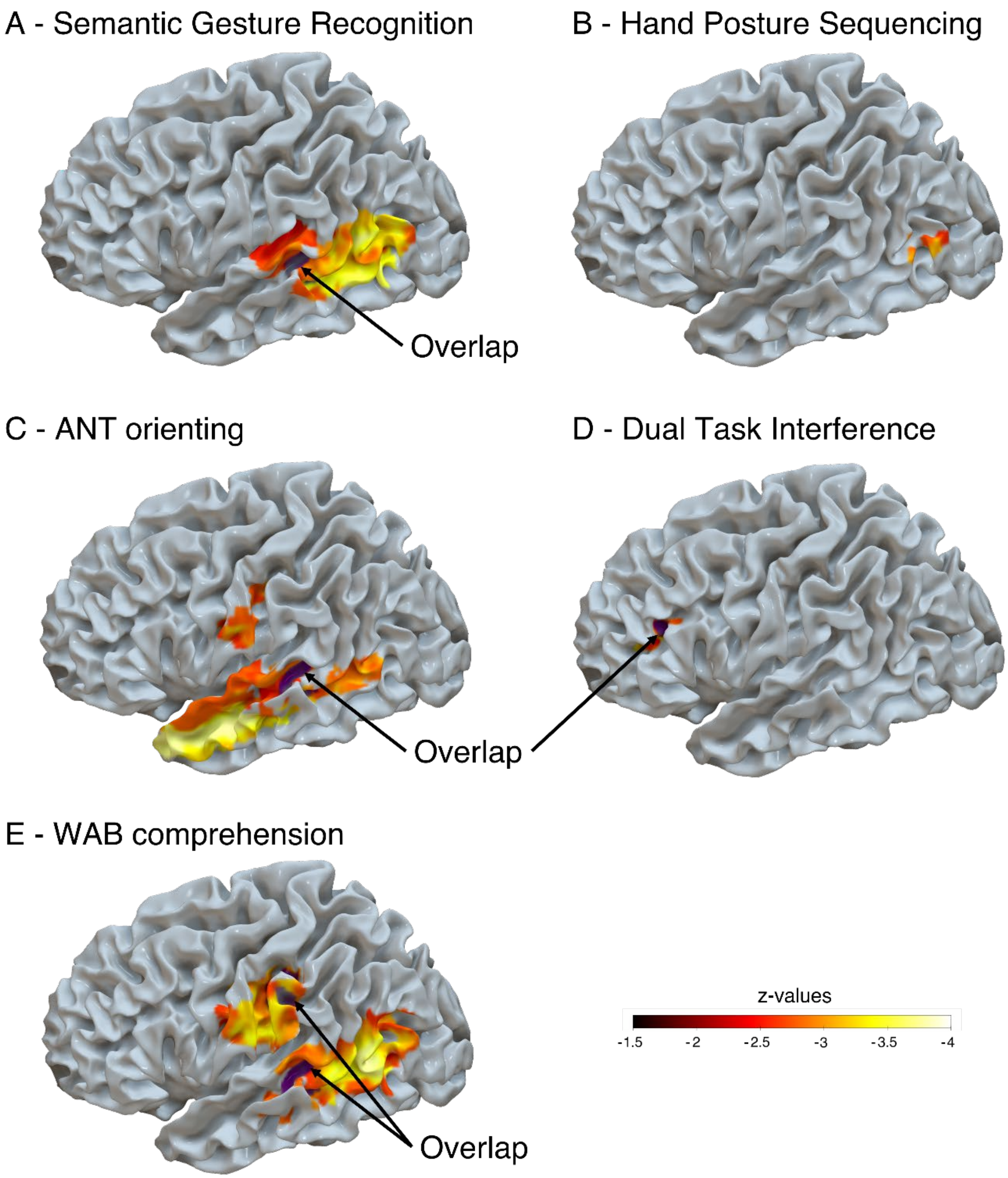


Figure S1: Lesion-symptom mapping analysis conducted for each of the tasks predicting NAT performance. Brain surface renders depict the voxels associated with impaired behavior for semantic gesture recognition (A), hand posture sequencing (B), ANT orienting score (C), dual-task interference (D) and WAB comprehension (E). Overlap in the voxels associated with impaired NAT performance are depicted in purple and indicated by black arrows. Maps of areas with significant voxels were thresholded at p-value < 0.005 one-sided, corresponding to a z-value < -2.57 and a cluster size of at least 100 voxels (i.e., 100 $mm^3$).
